## Supplementary Figure for "Rhythmic sampling of multiple decision alternatives in the human brain"

### Supplementary Information

#### Relationship between attentional saccades and pupillometry

We assessed how and if attentional saccades are related to neuromodulatory activity. Recent literature highlighted increasing pupil diameter as a peripheral measure associated with overtly switching attention[1,2]. We thus computed the pupil size as well as the temporal change in size in relationship to the likelihood of (covert) attentional saccades (Supplementary Fig. S8a,b). We identified a positive relationship of pupil diameter with the occurrence of stay events ($r_{mean}$ = 0.10, t19 = 3.26, p = 0.004; Supplementary Fig. S8c) as well as a larger pupil diameter during staying (p = 0.005; Supplementary Fig. S8d). Conversely, attention switches were negatively correlated with the change in pupil size ($r_{mean}$ = -0.07, $t_{19}$ = -2.87 p = 0.01; Supplementary Fig. S8c) and displayed event-triggered pupil constrictions (p = 0.01; Supplementary Fig. S8d). Our findings thus speak against an increase in pupil dilation during attentional switching. Rather, the pupillometric measures appear to display a general link between arousal and focused information sampling[3,4].

#### Supplementary Figures

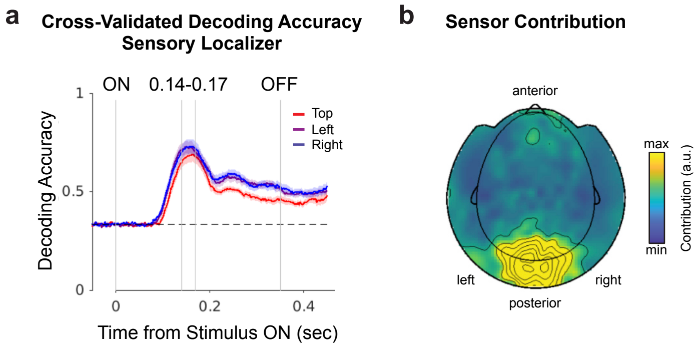


**Fig. S1 | Sensory localizer decoder performance and training time definition. a** Decoding accuracy for the attended target during the sensory localizer task over the stimulus presentation period. The decoder has been trained on all sensors and timepoints applying a leave one-block-out cross-validation approach (see Methods). The lines indicate the time-resolved accuracy values for the top (red), left (purple) and right (blue) target and the shaded areas indicate the SEM. The horizontal dashed line indicates chance level decoder performance. Vertical lines denote the stimulus onset, optimal decoder training period onset and offset as well as the stimulus offset (from left to right). **b** Sensor contributions to stimulus decoding during the optimal training period (highlighted in **a**). The contributions were derived as the summed principal component coefficients from the optimized weight matrix G (see Methods).

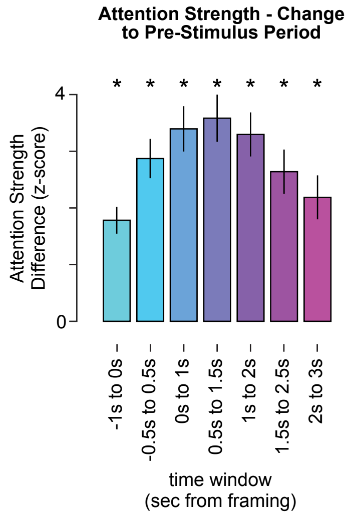


**Fig. S2 | Attention strength compared to pre-stimulus period.** Windowed and z-scored attention strength (vector length) over the stimulus presentation- compared to pre-stimulus period. Bars depict the average z-score difference for each stimulus period window (from early -1 to 0 s, light blue; to late 2 to 3s post-framing, purple; half overlapping windows) minus the pre-stimulus window (-1.5 to -1 s relative to framing cue). Vertical lines display the standard error of the mean (SEM) within each window. Asterisks denote significant differences ($p < 0.0001$). The t-scores from early to late windows are: $t_{19}$ = [7.72, 8.49, 8.71, 8.83, 8.70, 6.93, 5.79].

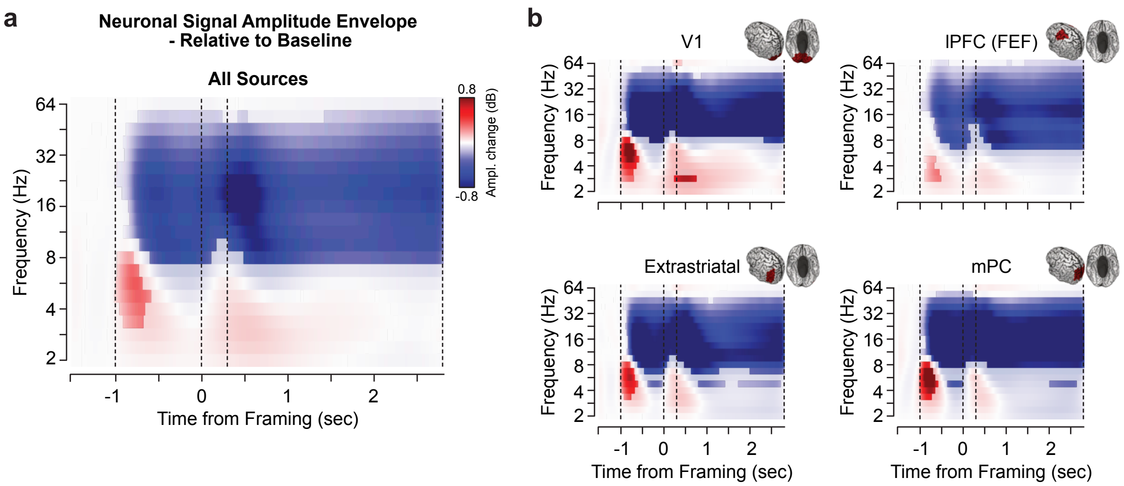


**Fig. S3 | Neuronal Signal Power Relative to Baseline. a,b** Time-frequency resolved distribution of the amplitude envelope for **a** all sources, and **b** subsets approximating early visual (V1, top left), extrastriatal and dorsal visual (bottom left), lateral prefrontal (lPFC with frontal eye-field FEF, top right) and medial parietal areas (mPC, bottom right). Red and blue colors indicate an increase or decrease relative to baseline, respectively. The values were statistically masked at $p_{FDR} < 0.01$ within each frequency. Vertical dashed lines denote the stimulus onset, framing on- and offset as well as the stimulus offset (from left to right).

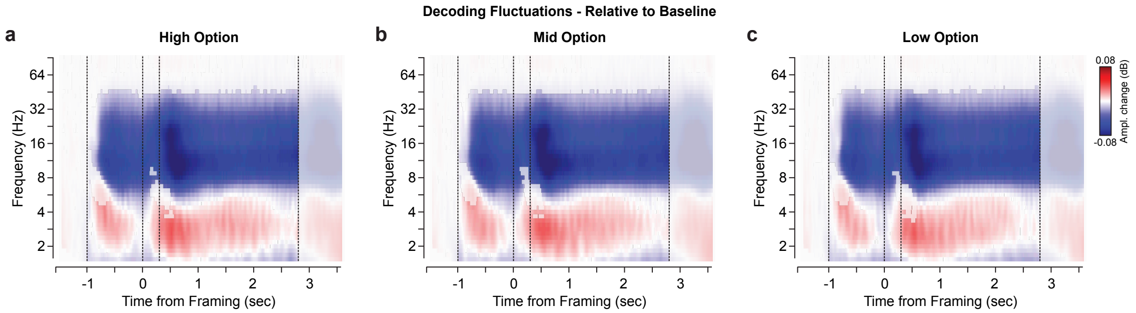


**Fig. S4 | Fluctuation of Stimulus Representation Decodability. a-c** Time-frequency resolved distribution of the amplitude of the decodability for the **a** highest, **b** second-highest and **c** lowest valued option. Red and blue colors indicate an increase or decrease relative to baseline, respectively. The values were statistically masked at $p_{FDR} < 0.01$ within each frequency. Vertical dashed lines denote the stimulus onset, framing on- and offset as well as the stimulus offset (from left to right).

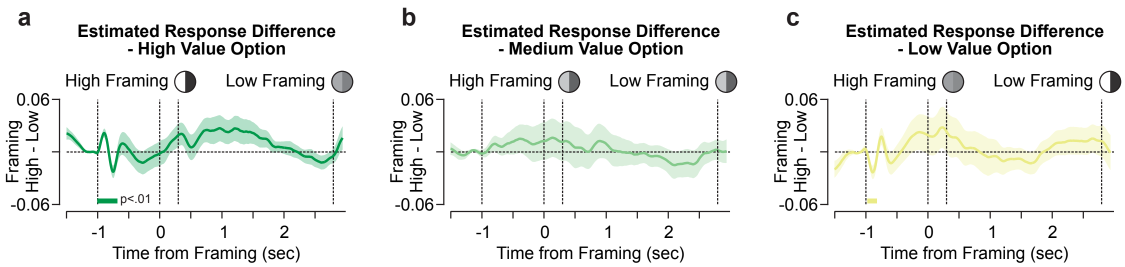


**Fig. S5 | Differences in estimated responses between framing conditions. a-c** Trial-averaged decodability (estimated response) differences for High- minus Low-Framing trials separately for **a** high, **b** medium, and **c** low decision value option within each respective trial. The circles schematically depict the compared contrast levels. Thick lines and shaded areas display the mean difference and the standard error, respectively.

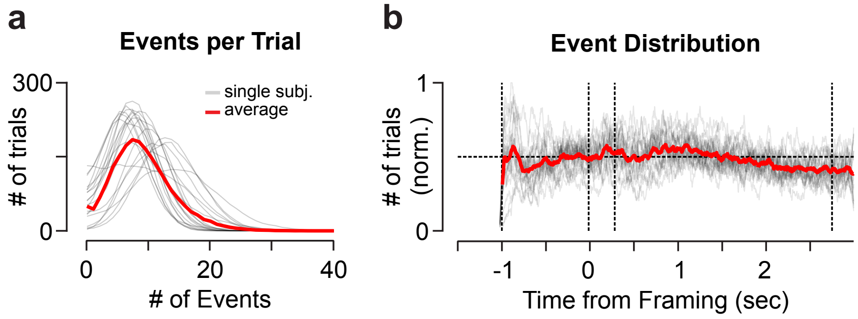


**Fig. S6 | Event characteristics according to the applied threshold. a** Histogram of the number of attentional event per trial and subject. The red line indicates the average over subjects, gray lines display the single subject distributions. **b** Distribution of events over the trial normalized within each subject (color-coded as in **a**).

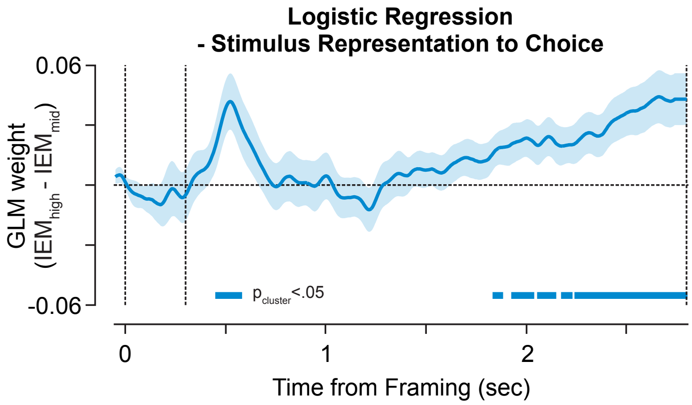


**Fig. S7 | Neural representation of decision alternatives predicts choices.** Time resolved logistic regression predicting choice from the relative decodability of decision alternatives ($IEM_{high}$ – $IEM_{mid}$), i.e. increased stimulus representation of the highest decision alternative, at each time point. The blue line and shaded area indicate the mean and SEM over participants. Thick lines at the bottom depict statistically significant differences from zero ($|t|_{19} > 0$, $p < 0.05$, cluster-mass permutation corrected). Dashed vertical lines denote the framing cue onset, offset and stimulus offset (from left to right).

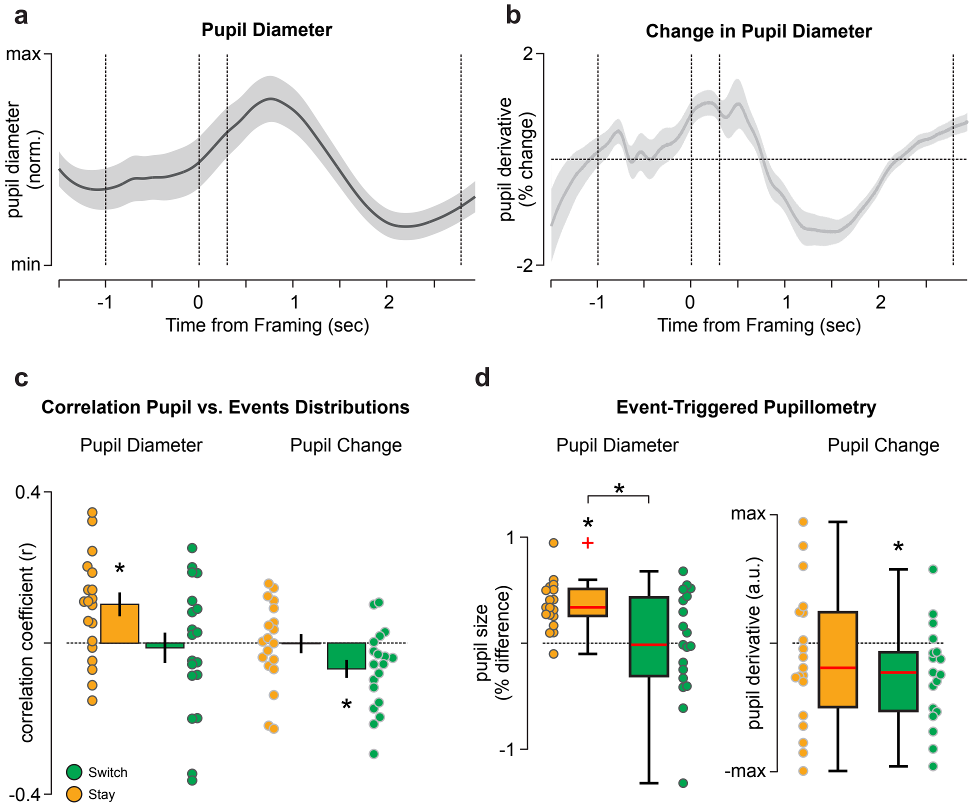


**Fig. S8 | Attentional saccade related pupillometry. a** The average normalized pupil diameter during task execution. The thick line indicates the average over subjects and the shaded area the SEM. Vertical dashed lines indicate the stimulus onset, framing cue onset and offset as well as the stimulus offset (from left to right). **b** the temporal derivative of the pupil size **a** normalized as percent-change. The thick line indicates the average over subjects and the shaded area the SEM. The vertical bars denote stimulus epochs (see above). **c** the correlation of the attentional event probability (Supplementary Fig. 5) with the average pupil size **a** and its temporal derivative **b** for each participant. The bars and vertical lines display the average and SEM over subjects, respectively. Orange and green indicate stay and switch attentional events, respectively. The colored dots represent the single subject Pearson correlation coefficients. Asterisks denote significant correlations ($|r| > 0$, $p_{FDR} < 0.05$). **d** box-plots of peri-event pupil size (left panel) and its temporal derivative (pupil change, right panel) as a function of attentional event types. The pupil size has been normalized against random events (see Methods). The colored dots display single subject values. Asterisks denote significant changes to random (pupil size) or against zero (change in pupil size, $p_{FDR} < 0.05$). Color-coded as above.

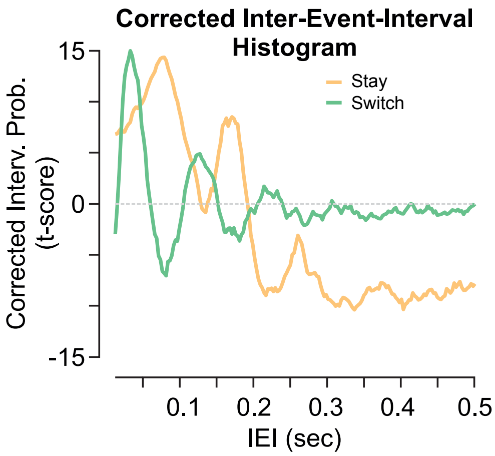


**Fig. S9 | Corrected inter-event-interval histogram.** Corrected inter-event-interval histograms separately for refocus and switch attentional events as t-score against random (compare Fig. 5). The horizontal dashed line indicates zero.

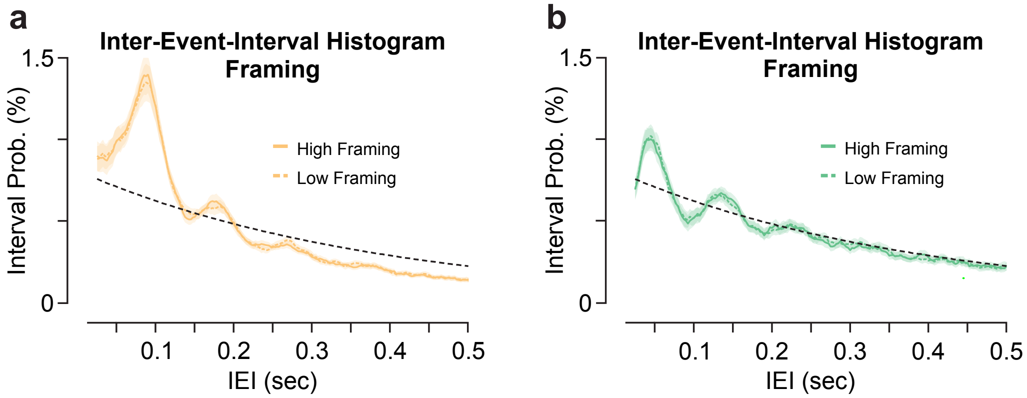


**Fig. S10 | Uncorrected inter-event-interval histograms separately for high and low framing trials. a,b** Uncorrected inter-event-interval histograms separately for **a** stay and **b** switch attentional events in High- (colored solid line) and Low-Framing conditions (colored dashed line; compare Fig. 5). The black dashed line indicates event intervals under random conditions.

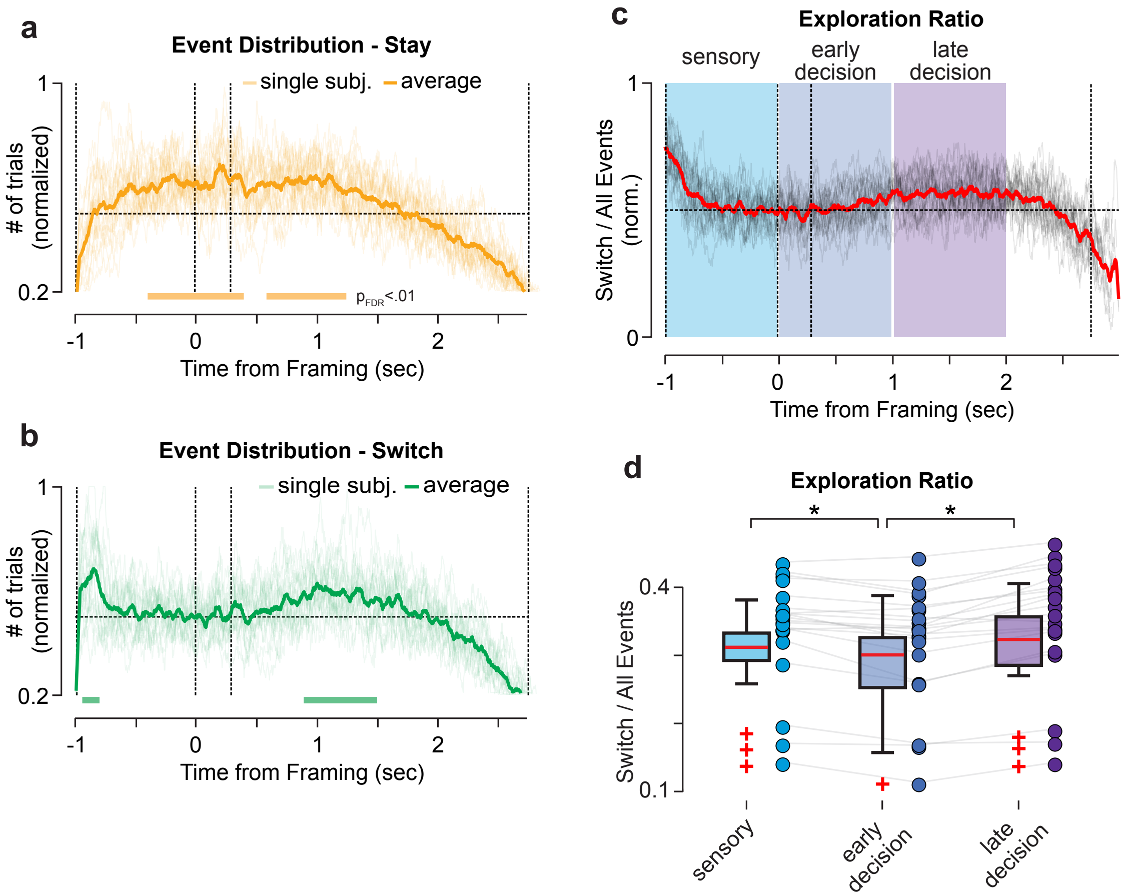


**Fig. S11 | Distribution of normalized event probability over the course of the trial. a-b** The distribution of event probabilities over the trial for **a** stay (orange) and **b** switch events (green). Transparent lines indicate normalized single-subject probabilities, opaque lines the average over subjects. The data was smoothed with a 0.05 seconds boxcar kernel for visualization. **c,d** Exploration ratio **c** over the course of the trial and **d** averaged over separate epochs of the trial. The exploration ratio describes at each time point the probability of an occurring event being a switch (data in **b** divided by (**a**+**b**)). The colored areas denote three non-overlapping epochs of the trial. Asterisks indicate significantly different exploration ratio between epochs (paired t-test, $p < 0.05$, Bonferroni corrected).

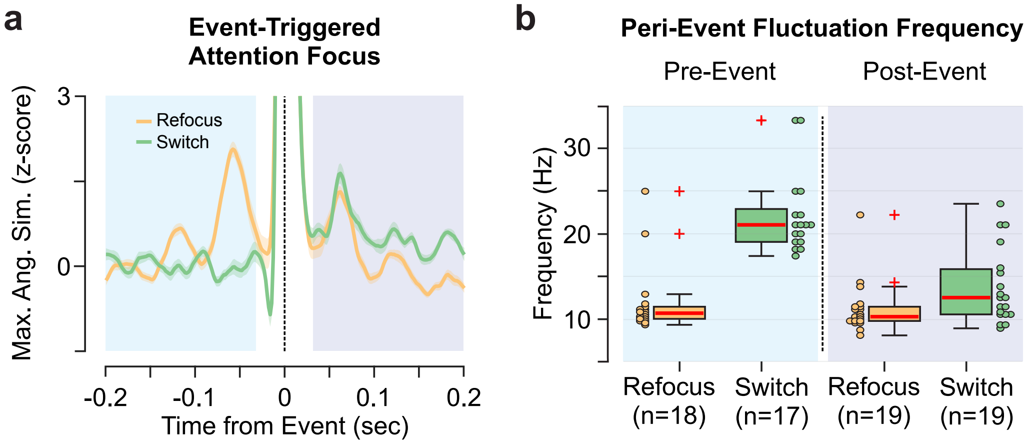


**Fig. S12 | Peri-event fluctuations of attention focus. a** Attentional event-triggered average of the attention focus, i.e. maximum angular similarity. The data was z-scored while excluding the center 0.06 seconds (event ± 0.03 sec.). The colored lines indicate the average over events and subjects separately for stay (orange) and switch events (green). The shaded areas indicate the SEM. The pre- and post-event epoch is highlighted in light blue and purple, respectively. **b** Dominant peri-event fluctuation frequencies defined via the average peak-to-peak latency within each epoch **a**. N denotes the number of subjects for which a peak-to-peak fit was possible.

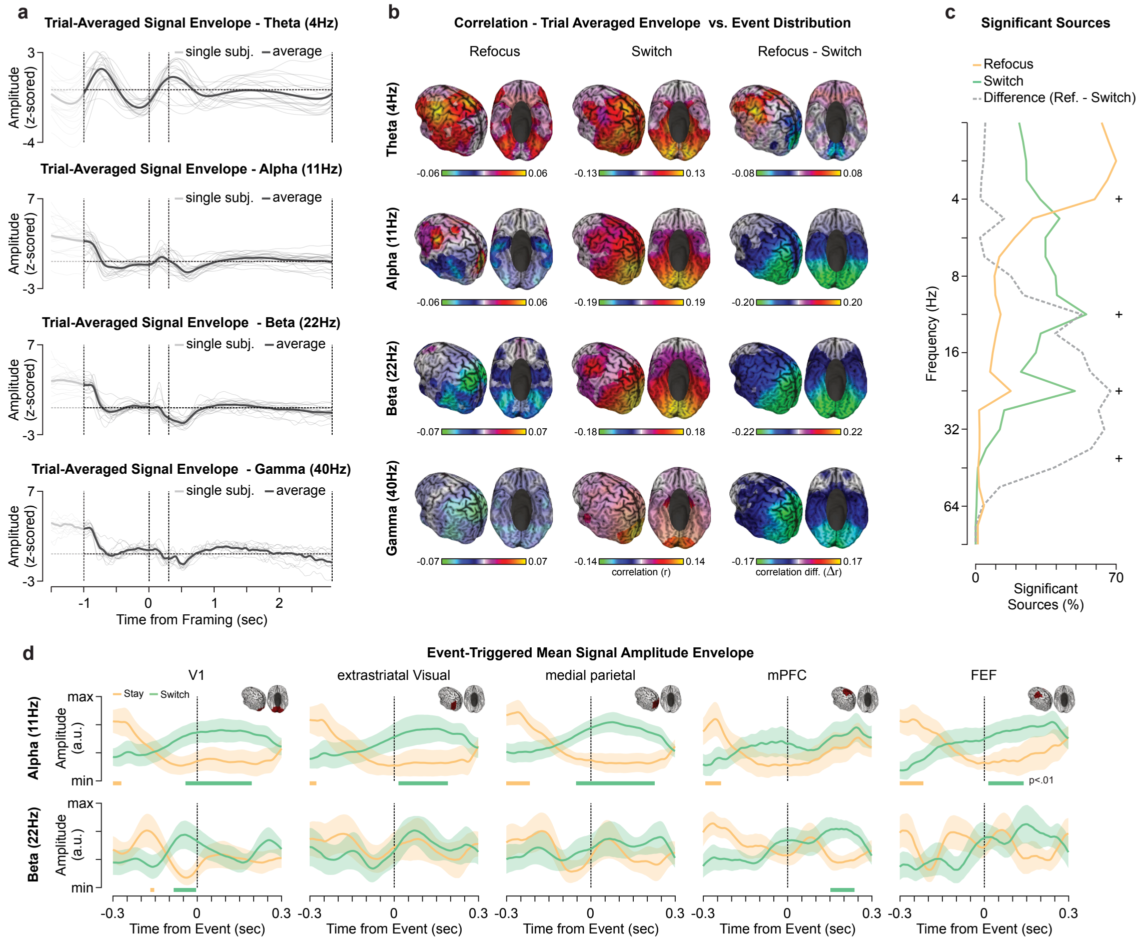


**Fig. S13 | Cortical dynamics of covert attentional events. a** Grand average of neuronal signal amplitude envelopes over sources, trials and participants. The values were z-scored within each subject prior to averaging. Transparent and opaque lines depict single-subjects and the average over subjects. From top to bottom the panels display the signal amplitudes for theta (4 Hz, Morlet wavelet with 0.5 octave bandwidth), alpha (11 Hz), beta (22 Hz), and gamma (40 Hz) carrier frequencies. **b** Cortical distribution of the correlation of **a** the trial averaged signal amplitude with the attentional event probability (Supplementary Fig. S5). Left and middle column show the correlation for stay and switch events, respectively. The right most column shows the difference. The color-scale was adjusted within each panel to ± the 98% percentile of the absolute value of correlations (|r|). Opaque sources indicate significant correlations ($p < 0.01$). The values were averaged over hemispheres for visualization. **c** Number of significant sources for the correlations in Figure S8B over the entire spectrum (2.8 - 64 Hz) separately for stay (orange), switch (green), and the difference (stay-switch; gray dashed line). Plus signs denote frequencies displayed in **b**. **d** Event-triggered signal amplitude within 5 regions of interest (early visual cortex (V1), extrastriatal visual cortex, medial parietal cortex, medial prefrontal cortex (mPFC), & dorsolateral PFC and frontal eye field (FEF); see inlays) in the alpha (11Hz) and beta (22Hz) frequency range. Green and orange lines depict the average activity around switching and staying attentional events. Shaded areas indicate the SEM. Thick bars denote significant differences between switch ($t_{Sw>Ref}$, green) and stay ($t_{Ref>Sw}$, orange; $p < 0.01$).

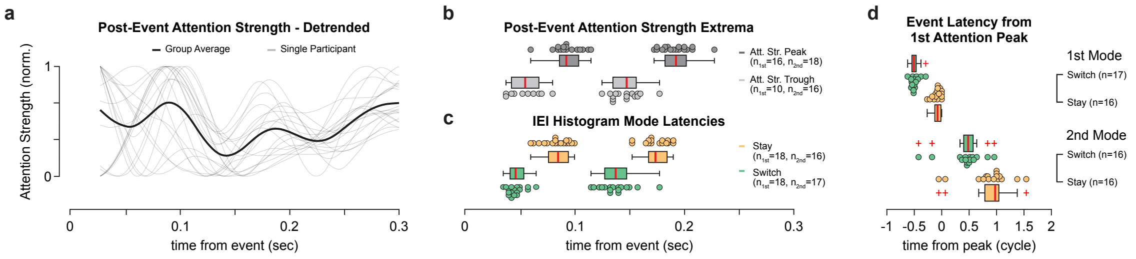


**Fig. S14 | Temporal relationship between post-event attention strength and attentional events. a** Post-event attention strength detrended and smoothed for each participant (grey lines) and the group average (black line). The signal was detrended within the displayed 0.3 s window and smoothed for each participant. **b** The box plots indicate the distribution of first (left) and second (right) peaks (top) and troughs (bottom) of the post-event attention strength over participants. Peaks were defined as local maxima, troughs as local minima preceding peaks on the single-participant level. **c** Absolute first- and second event mode latencies (in seconds) for switch (green) and stay (orange) events (see Fig. 5h for relative latency). **d** Relative latency between the first post-event attention strength peak and the first- and second event modes (color coded as above) in individual attention strength cycle length (see Fig. 7d).

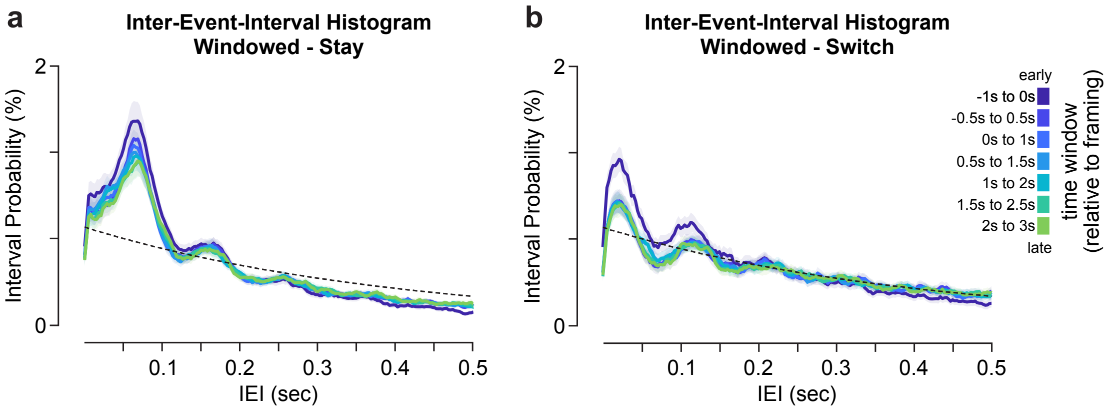


**Figure S15 | Inter-event-interval histograms within different epochs of the trial.** IEI histograms for **a** stay and **b** switch events color-coded from early (earliest -1s to 0s, relative to framing cue; blue) to late windows (latest 2s to 3s; green). The windows are half-overlapping and last 1 second each. Thick lines and shaded areas indicate the mean and standard error over participants.

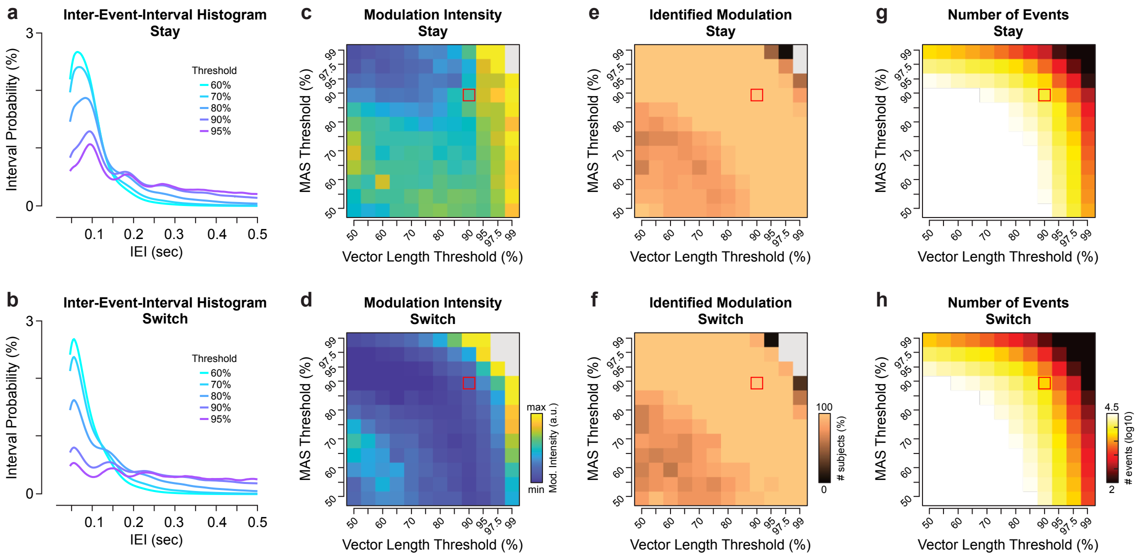


**Fig. S16 | Systematic assessment of threshold choices for the central findings. a,b** inter-event-interval histograms for different threshold choices separately for **a** stay and **b** switch events averaged over subjects. The percentile-thresholds were kept constant between the maximum cosine similarity (MAS) and attentional vector length at levels of 60%, 70%, 80%, 90% & 95% (see Methods). The lines were color-coded according to the threshold choice. **c,d** median modulation intensity between the first two peaks for the fully analyzed threshold space between 50% to 99% threshold varied separately for MAS and the vector length. The modulation depth defines the standard deviation in the IEI-range between the first- and second peak of the histogram (see Methods). Warmer colors indicate stronger modulation, i.e. more variable peaks and troughs. The top and bottom panels show the modulation intensity for **c** stay and **d** switch events. Threshold combinations that didn't yield at least one subject with two peaks are displayed in gray. Red squares indicate he threshold combination applied throughout the main text and results. **e,f** proportion of subjects with at least two nominal peaks separately for **e** stay and **f** switch attentional events. These subjects suffice the definition to compute the modulation intensity. Threshold combinations that didn't yield at least one subject with two peaks are displayed in gray. **g,h** the average numbers of identified events by threshold combination separately for **g** stay and **h** switch events.
